# Biophysical modeling of VIM to assess contributions of oscillatory activity to essential tremor

**DOI:** 10.1101/339846

**Authors:** Shane Lee, David J Segar, Wael F Asaad, Stephanie R Jones

**Affiliations:** Department of Neuroscience, Brown University; Carney Institute for Brain Science, Brown University; Alpert Medical School Department of Neurosurgery, Brown University; Brigham and Women’s Hospital Department of Neurosurgery; Norman Prince Neurosciences Institute at Rhode Island Hospital; Providence Veteran’s Affairs Medical Center, Center for Neurorestoration and Neurotechnology

## Abstract

Essential tremor (ET) is the most common movement disorder, in which the primary symptom is a prominent, involuntary 4–10 Hz rhythmic movement. The presence of tremor frequency oscillations (TFOs) in the ventral intermediate nucleus of the thalamus (VIM) is well-established, but it is often assumed that it is driven by cerebellar tremor frequency activity, while the role of intrinsic oscillatory activity in VIM is not clear. An improved understanding of the mechanisms of tremor and non-tremor frequency activity in VIM is critical to the development of improved pharmacological and neuromodulatory therapies. Starting from a canonical model of thalamus, we developed a biophysically-principled computational model of tremor field activity in the VIM, coupled with the thalamic reticular nucleus (TRN). We simulated TFOs in the model generated either by extrinsic tremor-periodic drive or intrinsic VIM-TRN interaction to understand whether these networks exhibited distinct biophysical properties, which may impact the efficacy of pharmacological or stimulation treatment for TFOs. Extrinsic and intrinsic TFOs in the model depended on T-type Ca^2+^ channels in different ways. Each also depended on GABA modulation in a site- and type-specific manner. These results suggested that efficacy of pharmacological manipulations may depend upon the mechanisms generating TFOs in VIM. Simulated non-tremor-related motor activity from cerebellum decreased extrinsic but increased intrinsic TFOs. Our results suggest that both mechanisms may be important to understand the emergence and cessation of TFOs in VIM and lead to experimentally testable predictions on how to modulate tremor frequency activity to improve treatment strategies for ET.

**Significance Statement:** Essential Tremor (ET) is a movement disorder in which the primary symptom is a prominent, involuntary, and rhythmic shaking, often of the hands. Electrical activity in many areas of the brain exhibit rhythmicity related to the patient’s tremor. One such area resides in a structure called the thalamus, but it is not fully known what gives rise to tremor-related activity. We created a computational model of this activity, which suggested how to differentiate tremor mechanisms and how these differences may contribute to other impairments in ET. Knowledge of the biophysical mechanisms contributing to tremor can ultimately lead to improvements in treatments to alleviate symptoms of ET.

## 1 Introduction

Essential Tremor (ET) is the most common movement disorder in adults, affecting 5-6% of the population [35, 36]. ET is dominated by a low frequency (4–10 Hz), involuntary “intention” tremor, often of the upper limbs [13, 36], but is no longer considered a monosymptomatic disorder [32], as both motor and cognitive symptoms have been observed [31, 37, 56]. Potential cerebellar dysfunction has been implicated in the etiology of ET [9, 13, 14, 37, 58], though the pathophysiology is heterogeneous and not fully understood [6, 17, 37, 48]. Tremor, by far the most dominant symptom, has been associated with tremor frequency oscillatory neural activity in the thalamus [25, 29, 52], primary motor cortex [20, 21], cerebellar cortex [52], and brainstem [52].

The origin—or origins—of tremor frequency oscillations related to tremor in ET are not fully known, though prior work has suggested that tremor frequency oscillations in the thalamus are driven by cerebellar activity [15, 37, 52]. It remains unclear though unlikely that there is a single driver of tremor frequency activity [6, 48], and the role of intrinsically generated tremor frequency activity within the thalamus is not fully known.

Pharmacologically, both T-type Ca^2+^ and *γ*-aminobutyric acid (GABA) activity has been implicated in the tremor pathophysiology in ET. Reduction of T-type Ca^2+^ channels in ET and other movement disorders has been suggested to reduce pathological oscillatory activity [19, 27, 50]. GABA-ergic modulation has been implicated in multiple aspects of tremorgenic oscillatory activity, particularly in the cerebellum [15, 18, 38, 52] but also within the VIM [1, 4, 15]. The “GABA hypothesis” suggests that disinhibition of pacemakers in the dentate nucleus of the cerebellum may lead to increased tremor frequency activity in the thalamus and cortex [15, 44], though other work suggests that the inferior olive generates and transmits tremor frequency activity to the cerebellum and then to the thalamus [14, 15, 18, 52]. Both T-type Ca^2+^ and GABA are modulated at multiple sites in the cerebellothalamocortical circuit, including the VIM, and how these individual sites contribute to the pathophysiology is not fully known.

An improved understanding of the cellular and circuit mechanisms generating tremor frequency neural activity in ET may lead to more effective pharmacological or electrical treatments. Prior computational modeling has investigated multiple aspects of the timing of deep brain stimulation on thalamic cells, including rebound bursting [3, 5, 32, 59], and recent work investigated mean field tremor frequency activity in thalamus [65]. Yet to our knowledge, within the VIM, whether tremor frequency neural activity is intrinsically or extrinsically modulated has not been fully established and is essential to understand mechanisms of generation toward improving treatment strategies.

Here, we developed a biophysically-principled computational model of tremor frequency activity in the VIM, based on a canonical model of thalamic oscillatory activity that involves reciprocal interaction with the thalamic reticular nucleus (TRN) [10].

We compared two distinct modes of generation of TFOs: extrinsically driven and intrinsically generated. Both T-type Ca^2+^ and GABA channels are broadly involved in oscillatory activity in the thalamus [10, 12], so these currents were varied in the model.

Our goal was to assess the extent to which intrinsic versus extrinsic mechanisms may contribute to tremor and how differences in these networks respond differently to pharmacological manipulations or afferent inputs representing non-tremor signals. In a companion paper, we investigated the differential impact of deep brain stimulation on TFOs produced by extrinsic or intrinsic mechanisms [33].

## 2 Methods

### 2.1 Overview of the model network

We sought to understand the biophysical cellular and circuit level contributions to tremor frequency neural activity in VIM. We used NEURON 7.5 [7, 23] to simulate a small volume of VIM—which might be considered a tremor cluster [46]—consisting of 25 multicompartmental thalamocortical VIM relay cells and 25 multicompartmental thalamic reticular nuclear (TRN) cells (Fig. 1). The basic network was based on prior modeling of networks of thalamocortical relay cells [12] and thalamic reticular network cells based on electrophysiological recording of ferret thalamus [10–12]. Bursting VIM cells, a prominent feature in ET [25], were also present in these models. Though the original work was a model of sleep or anesthesia, bursting modes underlying oscillatory activity have also been shown to be important in normal function [54], and we hypothesized that similar mechanisms may underlie pathological VIM bursting.

**Figure 1:**
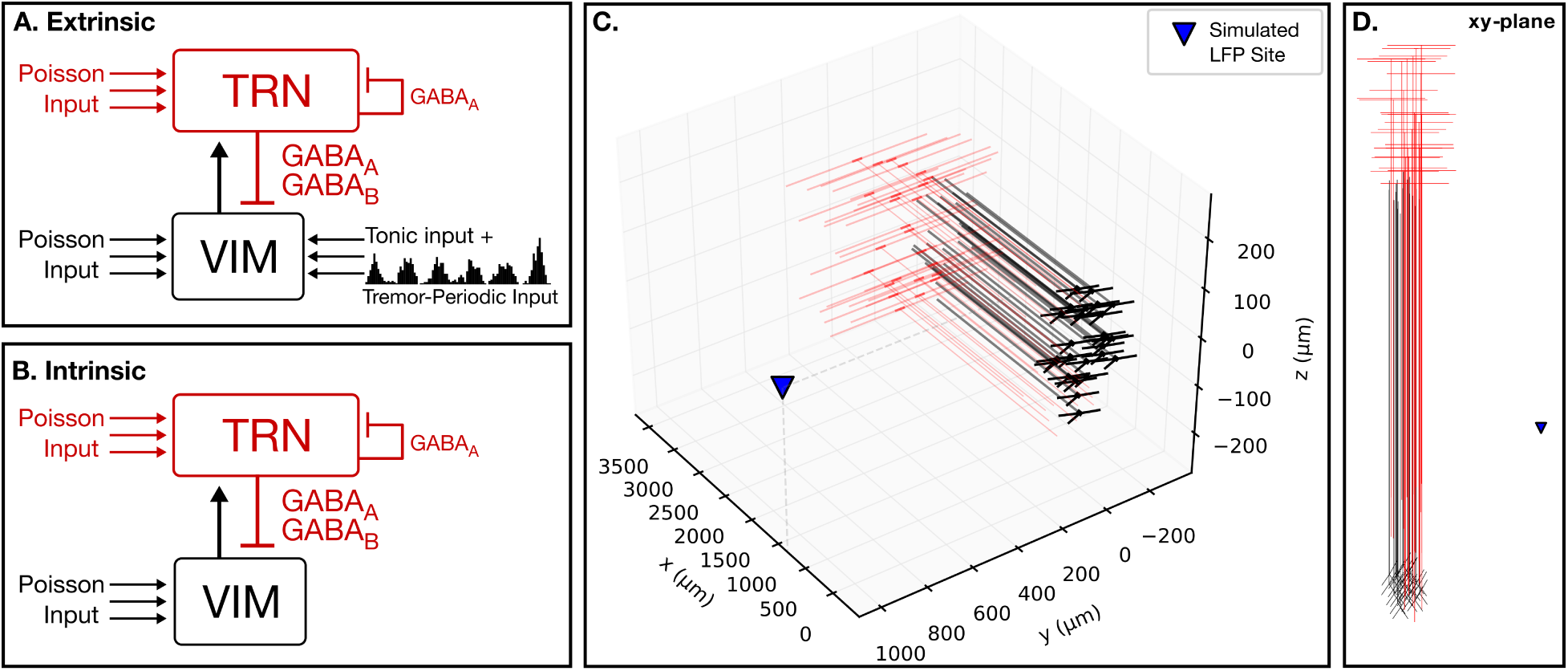
Model VIM-TRN networks. 25 multicompartment VIM cells (black) and 25 multicompartment TRN cells (red) were simulated. Background Poisson-distributed input was delivered to all cells. A. The “extrinsic” network generated tremor frequency oscillations through a combination of a phasic, tremor frequency synaptic input and tonic applied current input to the VIM cells. VIM cells made AMPA-ergic synaptic connections with the TRN cells, and TRN cells made GABA-ergic connections back to the VIM and also GABA_A_ connections within the TRN cells. B. The “intrinsic” network generated tremor frequency oscillations that were a product of glutamatergic excitation of TRN cells and reciprocal GABA-ergic inhibition of VIM cells, which caused post-inhibition rebound. This network was initiated by only the Poisson input to the VIM and TRN cells (see Methods). C. 3D perspective of network. Extracellular recording was simulated at (1250, 1000, 0) *μm* (blue triangle). D. xy-plane perspective. The center of mass of the VIM centered about the origin (0, 0, 0), with axons in the positive x direction toward the TRN. Both VIM and TRN cells in the networks received Poisson input.

We compared properties of the network for two possible mechanisms of tremor frequency oscillations: extrinsic and intrinsic (Fig. 1A–B). In both networks, VIM cells created AMPA-ergic synaptic connections with the TRN cells, and TRN cells created both GABA_A_ and GABA_B_ connections with the VIM cells. TRN-TRN GABA_A_ connections were also created reciprocally.

VIM cells (outlined in detail below) in the extrinsic model were driven by a tonic applied current of 0.06 nA to mimic the thalamic “tonic” mode that did not auto-generate oscillations [54] but responded to tremor-frequency AMPA-ergic synaptic inputs faithfully. A phasic input with a mean frequency of 6 Hz drove activity in this network. The input was initiated at 50 ms, and on each cycle of the input, 30 synaptic inputs were simulated with a Gaussian temporal distribution (*σ*=20 ms) and unitary weights of 4.25×10^−3^ *μS*.

In the intrinsic model, oscillations generated in the tremor frequency range were based on reciprocal interactions between the VIM and TRN cells. The intrinsic model had no tremor-frequency synaptic drive.

VIM cells were chosen from a uniform spherical distribution centered about the origin, with a cell density of approximately 15,000 cells per mm^3^ [24] (Fig. 1C–D). The volume of VIM cells simulated was approximately 2.5 mm away from the shell of TRN along an x-axis (arbitrarily chosen), based on the overall size of VIM estimated at 5 × 4 × (0.5–3) mm^3^ [16].

The 25 TRN cells (outlined in detail below) were assumed to be near the VIM cells, and their axons were projected directly into the volume of VIM. The TRN somata were also uniformly distributed but throughout a volume spanning the circumference of the VIM sphere in two dimensions, with a TRN depth of 1 mm (approximated from [27]).

### 2.2 Thalamocortical VIM Cells

#### 2.2.1 Geometry and Biophysics

The thalamocortical VIM cells were modeled using Hodgkin-Huxley dynamics after the 3-compartment cell from Destexhe [12]. In general, cell membrane voltages v_*i*_, for compartment *i* were given by the form of Eqn. 1, a sum of ionic, adjacent compartmental, synaptic (I_*syn*_), and external input (I_*external*_) currents, where *C* is the capacitance. Ohmic currents were a function of the compartmental voltage and the reversal potential E_rev_ for a particular ionic species.

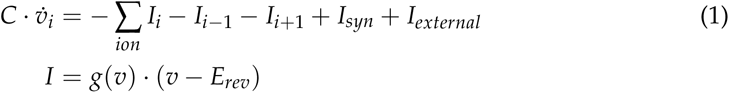

The original cell included a soma and a dendritic branch that consisted of two compartments. We adapted this model in several ways, summarized in Table 1A–B. First, we reduced the volume of the somatic compartment to conform to estimates of the size of human VIM cells [24]. Second, we included two more dendritic branches, radially outward from the soma, to reflect the stellate-like nature of these cells [24], and we included an axonal compartment. The VIM cells included all currents from the original model (Na^+^, K^+^, T-type Ca^2+^, and passive leak) and also included an additional potassium leak channel to sustain intrinsic T-type Ca^2+^-mediated oscillations in the VIM cells [10].

**Table 1:**
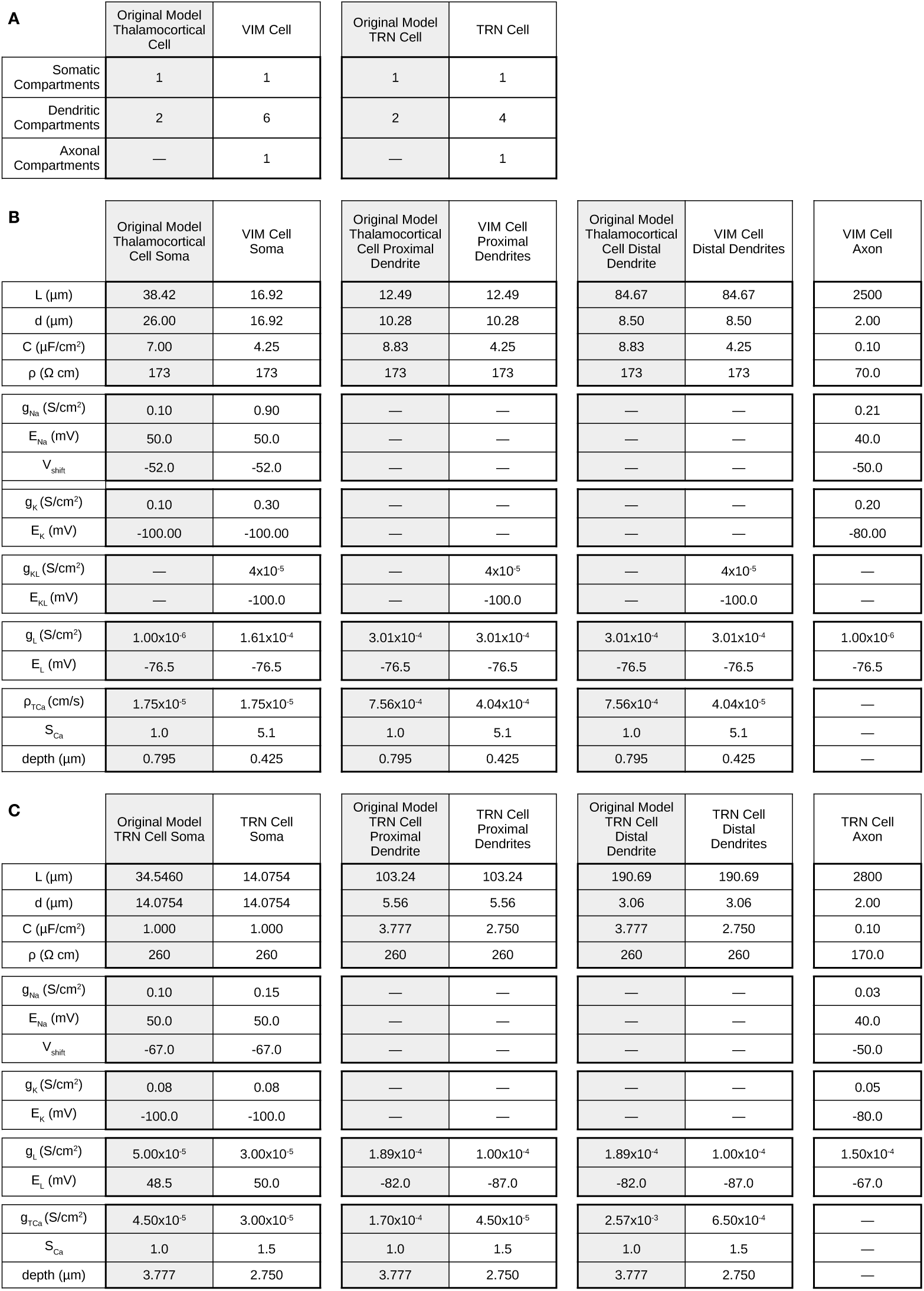
Key parameters compared to original model. Original model shaded in gray. A. Compartments in each section of the cells in the model. B. Geometry and biophysics from original model of thalamocortical cells [12] and the present VIM cell. C. Geometry and biophysics from the original model of TRN cells [10] and the present TRN cell.

The biophysics were subsequently tuned to mimic the dynamical properties of the original cell (Table 1B). In particular, the original thalamocortical cell somata had a cylindrical surface area of approximately 3138 *μm*^2^, larger than estimates in VIM of 500-900 *μm*^2^ [24]. Therefore we reduced the cylindrical surface area to 899 *μm*^2^.

We added two additional dendritic branches compared to the original model, totaling three dendritic branches, each consisting of two compartments. The dendrites were arranged in the xy-plane for simplicity but at different angles: *π*/4, 3*π*/4, and −3*π*/4 radians (see Fig. 1C–D). For each dendrite, the diameters and lengths from the original model were preserved (Table 1).

Axonal biophysical parameters were adapted from prior modeling work [41] and adapted to enable axonal spiking to follow somatic output. The axon was 2.5 mm in length, oriented along the +x axis, with a diameter of 2 *μm* and was unmyelinated for simplification, unlike previous work.

In some simulations, the T-type Ca^2+^ permeability (*ρTCa*, in the soma and dendrites) was varied for all compartments simultaneously by a scaling factor S_*Ca*_ that scaled the permeability of each compartment’s default value (listed in Table 1B). At baseline, S_*Ca*_ was 5.1. For simulations in which S_*Ca*_ was changed, values are reported relative to this baseline; for example, a relative value of 0.5 corresponds to a S_*Ca*_ value of 2.55.

#### 2.2.2 Noisy background inputs

For all VIM cells, a Poisson-distributed background synaptic input was also included into a distal dendritic compartment to give a baseline, non-tonic level of excitation to the network. The rate parameter *λ* was 10 ms, with a fixed strength of 5×10^−3^ *μS*. For all VIM and TRN cells, total number of segments in each compartment was set to 1 segment per 100 *μm*. One-third of cells also received −0.01 nA, due to estimates of the number of thalamic cells showing tremor frequency activity [25]. One-half of the remaining cells arbitrarily received 0.05 nA for heterogeneity, which did not make substantial changes in the spiking activity of the cells (data not shown).

#### 2.2.3 Dynamic properties of VIM cell compared to original reduced cell

The dynamical behavior of the original, two-compartment thalamocortical cell was preserved in the modified VIM cell (Fig. 2A). The original thalamocortical cell (Left panel of Fig. 2A) was simulated for 1.5 s, with the temperature parameter in NEURON set to 36°C. In the present VIM cell (Right panel of Fig. 2A), the initial membrane potential for all compartments was chosen from a Gaussian distribution with a mean of −85 mV and a standard deviation of 2 mV.

**Figure 2:**
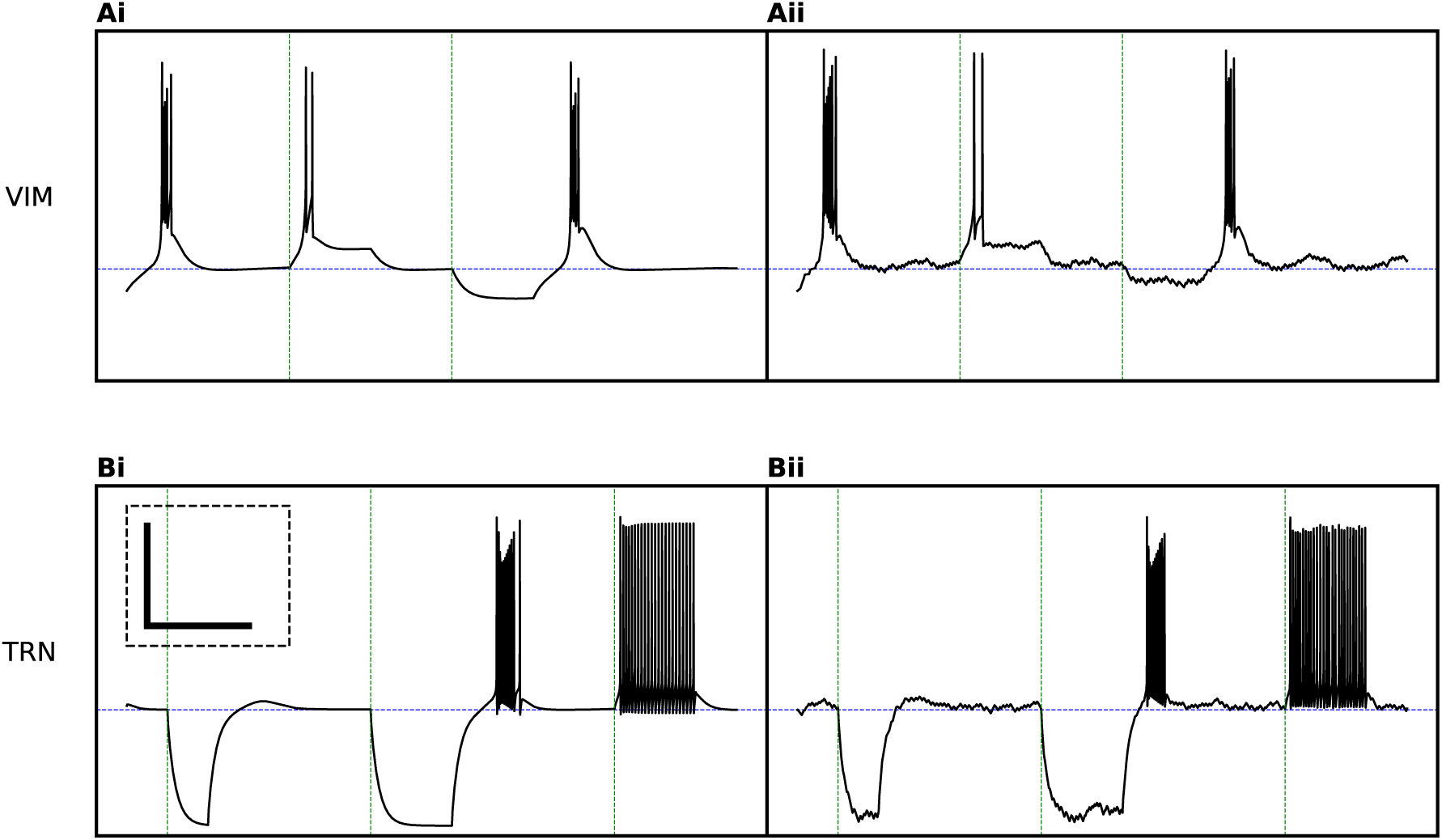
Model cells replicate original models. The VIM (Ai–ii) and TRN (Bi–ii) cells were modeled after Destexhe et al. [11, 12]. The original cells consisted of 3 compartments; here the VIM cells are 8 compartments (soma, 6 dendritic compartments, and an axon), and the TRN cells are 6 compartments (soma, 4 dendritic compartments, and an axon); see text for further details. For both cell types, 1.5 s are shown, the original cells are on the left (i), and the cells in the present model are on the right (ii). Poisson-distributed synaptic noise was applied to the new cells. A. For the VIM cells, the horizontal line is drawn at −74 mV. Both the original (Ai) and the new (Aii) cells were initiated at −80 mV, eliciting a burst of spikes. At 0.4 s (first green line), positive current was injected in the VIM cells for 0.2 s, giving rise to a doublet of spikes in each case. Finally, an inhibiting current was injected at 0.8 s for 0.2 s, and upon release, both VIM cells showed similar rebound spiking before returning to baseline. B. For the original (Bi) and new (Bii) TRN cells, the blue line marks −67 mV, the voltage at which the cells were initiated. At 0.1 s, for 0.1 s, a negative current was applied to the TRN cells, and no rebound spiking followed. However, at 0.6 s, the same negative amplitude current was applied for 0.2 s and elicited burst spiking in each of the cells. At 1.2 s, a positive current pulse of 0.2 s led to tonic spiking in both cells. Inset in original TRN cells applies to all plots; x scale is 0.25 s, and y scale is 50 mV.

Upon running the simulations, a short burst of spikes were elicited before returning to a steady state membrane potential of −74 mV. Two tonic current clamp stimulations were then applied. At 0.4 s (first green line in Fig. 2A), for 0.2 s, the stimulation amplitude was 0.07 nA in the original model and 0.17 nA in the VIM cell, which elicited two action potentials in each model. Upon release of the stimulation, the cells returned to baseline. At 0.8 s (second green line in Fig. 2A), for 0.2 s, the −0.10 nA were applied to both the original and new VIM cells. When this stimulation was released, a burst of action potentials mediated by T-type Ca^2+^ currents was elicited before the membrane potential returned to baseline.

### 2.3 Thalamic Reticular Nuclear Cells

#### 2.3.1 Geometry and Biophysics

The TRN cells were also modeled after three-compartment cells from Destexhe [11]. Similar to the VIM cell, we used the same generic formulation as Eqn. 1 and adapted the original TRN model in several ways. Though the soma and a dendritic branch were unchanged from the original TRN cell model, an additional dendritic branch—nearly identical to the original—was added to reflect lateral branching of cells through the TRN [47]. The dendritic arbor for each cell extended laterally within the model TRN, at opposite ends of the soma, at *π*/2 and −*π*/2, oriented within the TRN and perpendicular to the axonal projection to the VIM (see Fig. 1C–D). An unmyelinated axon was also added following prior work [41], with a length of 2.8 mm, oriented in the -x direction, and with a diameter of 2 *μm*. The same channels were inserted in the soma and dendritic sections: passive, Na^+^, K^+^, Ca^2+^ decay, and the original T-type Ca^2+^ channel [11]. A comparison of parameters is shown in Table 1A and 1C.

#### 2.3.2 Noisy Background Inputs

All TRN cells also had unique Poisson-distributed background AMPA-ergic synaptic inputs that contacted one distal dendritic compartment with a *λ* of 10 ms and a fixed strength of 5×10^−3^ *μS*, parameters identical to the inputs given to the VIM cells. One-quarter of cells randomly received 0.025 nA for heterogeneity.

#### 2.3.3 Dynamic properties of TRN cell compared to original reduced cell

Similar to the VIM cell, we simulated the original TRN cell under a variety of conditions and matched that behavior qualitatively (Fig. 2B). The original TRN cell (Left panel of Fig. 2B) and the new TRN cell (Right panel of Fig. 2B) were simulated for 1.5 s. The initial membrane potential for all compartments in the new TRN cell was chosen from a Gaussian distribution with a mean of −65 mV and a standard deviation of 2 mV. At 0.1 s (first green line in Fig. 2B), for 0.1 s, the stimulation amplitude was set to −0.4 nA in the original cell and −0.68 nA in the new TRN cell. Release of the current clamp did not result in rebound depolarization. After the membrane potential stabilized, at 0.6 s, the cell was clamped at the same amplitude for twice as long—0.2 s, which resulted in a burst of spikes following the release. At 1.2 s, a positive current of 0.1 nA to the original cell and 0.15 nA to the new TRN cell resulted in tonic spiking for the entire 0.2 s. The cells returned to a baseline of −67 mV.

### 2.4 Synapses

Following the original network model of thalamus [10], the following synapses were included in this model: fast excitatory AMPA-ergic from VIM to TRN, fast and slow inhibition from TRN to VIM, and fast inhibition from TRN to TRN. While VIM-TRN inputs originated at the distal axonal compartments, TRN to TRN connections were modeled without spatial specificity, using somatic presynaptic voltages that terminated on somatic compartments.

Fast inhibition approximated GABA_A_ in the model, while slow inhibition was a model for GABA_B_. As a simplification from the prior model, synaptic currents were modeled with a two-state kinetic scheme of summed exponentials (Eqn. 2–3) [7], were created all-to-all, and there were no autapses.

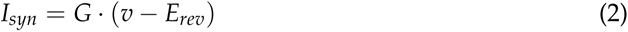

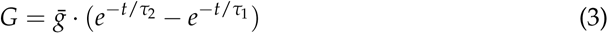

Conductances between VIM and TRN synaptic connections were set to generate tremor frequency activity in the intrinsic network (all units are μS): VIM-TRN AMPA conductance was 3.75×10^−4^; the TRN-VIM GABA_A_ conductance was 2.5×10^−5^, the TRN-VIM GABA_B_ conductance was 1.5×10^−4^, the TRN-TRN GABA_A_ conductance was 5×10^−4^. These conductance parameters were the same for both networks.

The excitatory AMPA receptors had a reversal potential of 0 mV, an exponential rise time *τ*_1_ of 0.5 ms, and an exponential decay time *τ*_2_ of 2 ms. GABA_A_ connections were modeled with *τ*_1_ of 0.5 ms, *τ*_2_ of 5 ms [28], and a Cl^−^ reversal potential of −85 mV. GABA_B_ connections had a *τ*_1_ of 1 ms, a *τ*_2_ of 100 ms, and a K^+^ reversal potential of −95 mV [10].

### 2.5 Additional Inputs

Aside from tonic currents and Poisson-distributed inputs applied to individual cells in the model (see above), externally sourced synaptic inputs were also simulated. To model ongoing, non-tremor inputs that might arise from cerebellum or cortex [32], we simulated additional Poisson-distributed inputs that terminated on AMPA-ergic synapses on the dendrites of the VIM cells. Each event time of the Poisson input consisted of 25 Gaussian-distributed inputs with a 3 ms standard deviation. The mean Poisson rate, at 40 sp/s, was within observed estimates of tonic spiking activity in the dentate nucleus [61]. The amplitude of the unitary input conductance was varied.

For the intrinsic network, we also simulated the effect of an external tremor-frequency input, modeled with an AMPA-like synaptic input that had an mean interevent interval of ~167 ms (6 Hz). On each cycle of the input, 30 Gaussian-distributed inputs arrived at the distal end of a dendritic branch with a standard deviation of 20 ms. The amplitude of this conductance was also varied.

### 2.6 Model Experimental Design and Statistical Analysis

A step size of 0.025 ms was used for all simulations, and 10 iterations were run for comparison across noise conditions. Sources of noise in the model included the specific realizations of the uniformly-distributed spatial locations of the somata for both the VIM and TRN cells, the Gaussian initial membrane potentials, and the Poisson inputs. For most comparisons unless otherwise noted, a non-parametric Mann-Whitney U test was used to assess significance [39, 62], which for all simulations was defined as *p* < 0.025 (n=10 simulations for all comparisons unless otherwise noted). Analyses were performed in Python 3.5 with standard modules, SciPy 0.18, and NumPy 1.13.

We were primarily interested in how tremor periodic activity in VIM would transfer to downstream areas such as primary motor cortex, so the principal output analyzed in the model was the axonal output spiking from the VIM cells. Individual spikes were identified as the membrane potential crossed a 0 mV threshold. Spike times from all VIM cells were concatenated, and the aggregate spike output was convolved with a Gaussian window with a width of 10 ms and a standard deviation of 2 ms. Mean VIM spike outputs were calculated among different simulations. All error is represented as standard error, with the number of samples specified.

Local field potentials were also simulated, using the extracellular mechanism in NEURON. The virtual recording electrode was placed in the volume at coordinates (1250 *μm*, 1000 *μm*, 0 *μm*), as seen in Fig. 1C–D.

Spectral analysis was performed on the convolved spike output and also on the simulated field potential, using a standard discrete Fourier transform in which the frequency domain signal was represented as a density of the total signal energy. Tremor energy was calculated as the concentration of total energy in the tremor frequency band (4–10 Hz), enabling comparisons across conditions.

For analyses comparing extrinsic to intrinsic tremor power, all values were directly compared across conditions for statistical tests. Some values were reported relative to a baseline condition for convenience, which was established by taking the mean of the baseline condition and normalizing all compared values to that value. Statistical tests were performed on raw values.

#### Code accessibility

The model is available on ModelDB (https://senselab.med.yale.edu/ModelDB/) with accession number 240113.

## 3 Results

### 3.1 Dual mechanisms in VIM-TRN model led to tremor frequency activity

Tremor frequency spiking activity and field activity are commonly observed features in the VIM [25, 26, 45, 53]. Yet it is not clear whether tremor frequency activity arises from extrinsically driven or intrinsically generated mechanisms, and a greater understanding of the origins of neural activity related to tremor may help to improve treatment for ET.

We sought to understand the role of cellular and circuit biophysics in the generation of tremor frequency local field potentials in the canonical thalamic circuit. We also aimed to simulate how different mechanisms of tremor frequency activity were affected by non-tremor-related motor activity. Though it is suggested that extrinsic cerebellar inputs drive tremor in thalamus [15, 37, 52], the intrinsic oscillatory capability of the thalamus is also well-described [10, 54]. We reasoned that the biophysical mechanisms underlying the intrinsic bursting mode of VIM may influence how tremor frequency activity is represented, which has implications for reduction of tremor. We modeled both extrinsic and intrinsic mechanisms of tremor frequency oscillations in the thalamus (Fig. 1A–B). Complete details of the networks can be found in the Methods.

Both the extrinsic and the intrinsic models consisted of reciprocally connected ventral intermediate nucleus of the thalamus (VIM) and the inhibitory thalamic reticular nucleus (TRN) that exhibited TFOs. All simulations included 25 VIM cells, coupled all-to-all to 25 TRN cells. An example schematic geometry of the network is shown in Fig. 1C–D. Individual VIM and TRN cells in the model were modified from prior work [10–12] by adding compartments and tuning the biophysics to exhibit similar behavior under a range of conditions (Fig. 2). We extended the original models with additional axonal and dendritic compartments in particular to be suitable for simulations of extracellular electrical “deep brain” stimulation, a widely-used therapy for medication-refractory ET, which we addressed in a companion study [33]. We analyzed spike rate outputs at the distal axon terminal of the VIM cells to understand how tremor frequency activity in VIM might be transmitted to downstream areas such as primary motor cortex and also simulated the extracellular field potential.

The extrinsic 50-cell model exhibited tremor frequency activity generated with a tremor-periodic AMPA-like synaptic input into tonically activated VIM cells (Fig. 1A and Fig. 3A–E). Here the peak frequency occurred at 6.4 Hz, and the tremor frequency concentration was 0.571 ± 0.018 mV^2^/Hz. The intrinsic 50-cell model was generated through the reciprocal interaction of GABA-ergic inhibition from TRN and T-type Ca^2+^ currents in the VIM cells, with a peak frequency of 6.7 Hz and a tremor frequency concentration of 0.572 ± 0.032 *mV*^2^/Hz (Fig. 1B and Fig. 3F–J). The tremor frequency power for the extrinsic and intrinsic models were not significantly different (*p* = 0.45).

**Figure 3:**
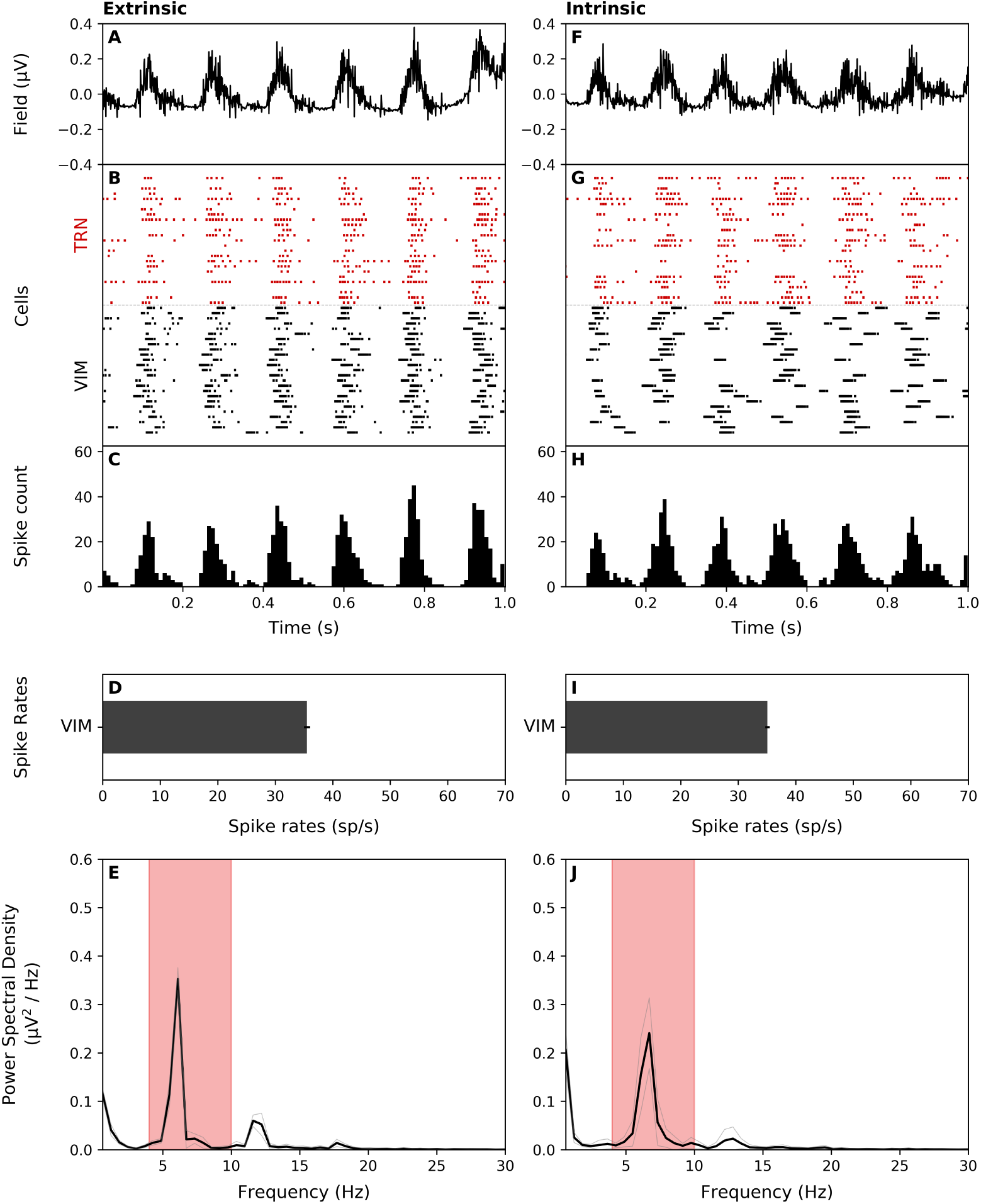
Extrinsic and intrinsic models of tremor frequency oscillations showed similar basic properties. A–E. Extrinsic tremor frequency oscillations were generated from tremor periodic synaptic drive. Data shown are simulated extracellular field (A), cell spike rasters for VIM (black) and TRN (red) cells (B), spike count histogram for VIM cells (C), mean VIM cell spike rates for all simulations (D), and mean power spectrum of spike count convolution for all simulations, with tremor spectral region highlighted in red (E). F–J. Same data for intrinsic oscillations that relied on VIM-TRN interactions. Mean VIM spike rates and tremor spectral concentration were not significantly different in these models. Spectral power at tremor frequencies (4–10 Hz) were also not significantly different.

VIM cell spike rates for both the extrinsic (35.5 ± 0.5 spikes/s) and intrinsic models (35.1 ± 0.39 spikes/s) were not significantly different (*p* = 0.29, Fig. 3D, I). Both were slightly higher than observed rates of 26.8 ± 5.3 spikes/s reported during behavioral tremor [26].

Overall, activity generated by the extrinsic and intrinsic models exhibited visible tremor frequency oscillations with similar statistical properties (Fig. 3).

### 3.2 Tremor frequency activity depended on T-type Ca^2+^ in VIM cells

Prior work in the thalamus has shown the importance T-type Ca^2+^ channels for pacing normally occurring oscillatory activity [10], and the function of T-type Ca^2+^ activity can be manipulated pharmacologically. The efficacy of treatments targeting T-type Ca^2+^ activity will therefore depend on the role of T-type Ca^2+^ channels on tremor frequency activity, and we hypothesized that these channels may also be important in sustaining pathological, tremor frequency activity. We varied the T-type Ca^2+^ conductance simultaneously in multiple compartments in VIM cells using a scale factor (S_*Ca*_) in both the extrinsic and intrinsic models and analyzed changes both in frequency and power (Fig. 4). Changes to S_*Ca*_ only affected the frequency of tremor frequency activity in the intrinsic but not the extrinsic network.

**Figure 4:**
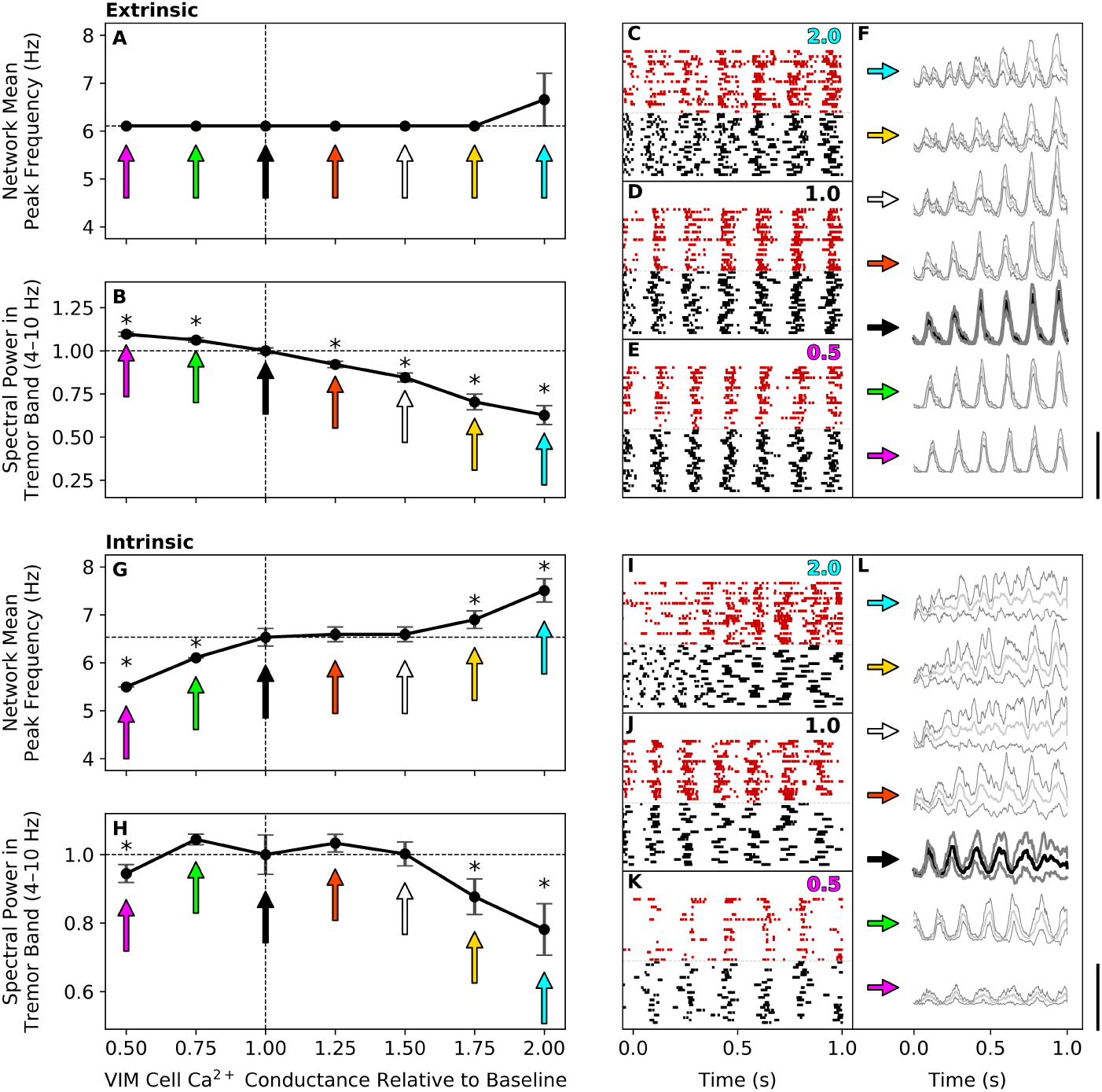
Changes in T-type Ca^2+^ conductance affect both extrinsic and intrinsic tremor frequency activity. T-type Ca^2+^ conductance changes changed spectral power in both networks but affected frequency only for the intrinsic network. S_*Ca*_ at baseline (1.0) was 5.1 (see Methods), and changes affected all T-type Ca^2+^ conductances in VIM cells but not TRN cells. Scale bar for F and L is 25 sp/s. All comparisons to baseline at 1.00. Values in Table 2. A–F. Extrinsic model. A. Changes in S_*Ca*_ did not affect the peak frequency significantly. B. Reductions in S_*Ca*_ increased tremor frequency activity, as the extrinsic input was the sole, rhythmic driver to the network. S_*Ca*_ increases reduced tremor frequency activity, as the temporal dynamics of T-type Ca^2+^ were strained. C–E. VIM (black) and TRN (red) spike rasters for examples at 2.0 (C), 1.0 (D), and 0.5 (E). F. Averaged spike convolutions with standard error for each conductance level; colors correspond to A and B. G–K. Intrinsic model. G. Increases in S_*Ca*_ increased the peak frequency but still within the tremor frequency range, and S_*Ca*_ decreases reduced the peak frequency. T-type Ca^2+^ dynamics influenced the properties of tremor frequency activity in the absence of an external periodic driver. H. Increases in S_*Ca*_ reduced power in the tremor frequency band, as in the extrinsic network, but in contrast, decreases also led to a significant reduction. In the absence of an external driver, the intrinsic model required T-type Ca^2+^ dynamics to sustain the strength of tremor frequency activity. I–K. VIM (black) and TRN (red) spike rasters for examples at 2.0 (I), 1.0 (J), and 0.5 (K). L. Averaged spike convolutions with standard error for each conductance level; colors correspond to G and H.

In the extrinsic model (Fig. 4A–F), neither reductions nor increases in S_*Ca*_ significantly affected the peak frequency of 6.1 Hz (Fig. 4A and Table 2). However, reductions in S_*Ca*_ led to significant increases in the tremor frequency band power, while S_*Ca*_ increases significantly reduced the power, for all levels tested, demonstrating a strong dependence of the tremor frequency power but not the peak frequency on S_*Ca*_ in the extrinsic model (Fig. 4B and Table 2).

**Table 2:** Modulation of tremor frequency activity by T-type Ca^2+^Conductance in models.

A. Extrinsic Model
| Relative<br>T-Type $\text{Ca}^{2+}$<br>Conductance | 0.50 | 0.75 | 1.0<br>($S_{\text{Ca}} = 5.1$) | 1.25 | 1.50 | 1.75 | 2.00 |
| --- | --- | --- | --- | --- | --- | --- | --- |
| Peak<br>Frequency<br>(Hz) | 6.10±0.00<br>p=0.500 | 6.10±0.00<br>p=0.500 | 6.10±0.00 | 6.10±0.00<br>p=0.500 | 6.10±0.00<br>p=0.500 | 6.10±0.00<br>p=0.500 | 6.65±0.55<br>p=0.184 |
| Tremor<br>Frequency<br>Power | *<br>1.098±0.036<br>p=3.8×10<sup>-4</sup> | *<br>1.063±0.038<br>p=0.004 | 1.000±0.051<br>(0.573±0.029 mV<sup>2</sup>/Hz) | *<br>0.921±0.056<br>p=0.023 | *<br>0.845±0.078<br>p=2.91×10<sup>-4</sup> | *<br>0.704±0.137<br>p=9.13×10<sup>-5</sup> | *<br>0.627±0.165<br>p=9.13×10<sup>-5</sup> |

B. Intrinsic Model
| Relative<br>T-Type $\text{Ca}^{2+}$<br>Conductance | 0.50 | 0.75 | 1.0<br>($S_{\text{Ca}} = 5.1$) | 1.25 | 1.50 | 1.75 | 2.00 |
| --- | --- | --- | --- | --- | --- | --- | --- |
| Peak<br>Frequency<br>(Hz) | *<br>5.49±0.00<br>p=2.55×10<sup>-5</sup> | *<br>6.10±0.00<br>p=7.14×10<sup>-3</sup> | 6.53±0.18 | 6.59±0.15<br>p=0.326 | 6.59±0.15<br>p=0.326 | 6.90±0.18<br>p=0.046 | *<br>7.51±0.24<br>p=0.002 |
| Tremor<br>Frequency<br>Power | *<br>0.945±0.079<br>p=0.013 | 1.045±0.047<br>p=0.425 | 1.000±0.172<br>(0.575±0.033 mV<sup>2</sup>/Hz) | 1.034±0.078<br>p=0.455 | 1.002±0.105<br>p=0.339 | *<br>0.877±0.157<br>p=0.023 | *<br>0.782±0.226<br>p=0.006 |

Raster data (Fig. 4C–E), as well as the spike histograms (Fig. 4F) illustrated how S_*Ca*_ changes affected VIM cells in the extrinsic network. With reduced S_*Ca*_ in the extrinsic network (Fig. 4E), the spike rasters showed stronger synchrony across the cells, as the extrinsic tremor drive had a greater contribution to the waveform width on each cycle of the oscillation. The reverse occurred with increased S_*Ca*_: the T-type Ca^2+^ dynamics began to interfere with the strong extrinsic drive, leading to a partial breakdown in synchronous activity in the tremor frequency band.

In contrast to the extrinsic model, S_*Ca*_ changes significantly affected the peak frequency of the intrinsic network (Fig. 4G–L and Table 2). Frequencies are reported relative to the baseline frequency of 6.5 Hz (“1.00”) with no changes in S_*Ca*_. Both reductions tested significantly lowered the peak frequency, both staying within the tremor frequency band. In contrast, increases in S_*Ca*_ significantly increased the peak frequency, also staying within the tremor frequency band. These differences suggested that if blocking T-type Ca^2+^ channels altered the frequency of tremor frequency activity in VIM, intrinsic mechanisms would be implicated, while the absence of frequency changes would be more consistent with extrinsic drive of frequency tremor activity.

At large deviations from the baseline S_*Ca*_ in the intrinsic model, significant changes in power occurred. Increases in S_*Ca*_ led to reductions of power in the tremor band, similar to that seen in the extrinsic network. However, S_*Ca*_ reduction by half led to a significant decline in tremor frequency power in the intrinsic network, opposite the effect seen for the extrinsic network. This suggests that, depending on the biophysical mechanism for tremor frequency oscillations in the VIM, blocking T-type Ca^2+^ channels would have opposite effects. However, tremor frequency activity in both networks was paradoxically reduced by increases in T-type Ca^2+^, suggesting that T-type Ca^2+^ agonists such as ST101 or SAK3 [55, 64] may work to reduce tremor in either case.

The raster data (Fig. 4I–K) and spike histograms (Fig. 4L) for the intrinsic network demonstrated different mechanisms of reduction of TFOs compared to the extrinsic network. The reduction in tremor frequency activity in the intrinsic network was due to the fact that there was no external driver to support the oscillations, which were driven by the response of T-type Ca^2+^ to the phasic inhibition from TRN. Increases in S_*Ca*_ in the intrinsic model also partially broke down oscillatory activity, despite a simultaneous increase in the frequency (Fig. 4I).

Taken together, changes in S_*Ca*_ affected both extrinsic and intrinsic tremor frequency oscillations in different ways, and the model results suggested that pharmacological manipulations of T-type Ca^2+^ activity directly in VIM lead to different outcomes. The model results predict that T-type Ca^2+^ antagonists would increase extrinsic tremor frequency activity and decrease intrinsic tremor frequency activity, while T-type Ca^2+^ agonists would reduce tremor frequency activity in either. Targeted experiments manipulating T-type Ca^2+^ in the VIM could help to disambiguate the mechanisms underlying tremor frequency activity in VIM.

### 3.3 Tremor frequency power depended upon GABA_A_ within the TRN but not from TRN to VIM

Tremor in ET is thought to involve GABA-ergic mechanisms [15, 22, 44], and medications that reduce tremor may also potentiate GABA [15]. However, GABAergic mechanisms exist both within the TRN and VIM, and the site of modulation and type of receptor may have a profound influence on the presence of tremor frequency activity. To test the effects of site-and type-specific GABA in the models, we first modulated GABA_A_ conductances at two different sites (illustrated in Fig. 1A–B): from TRN to VIM and from TRN to TRN, shown in Fig. 5 and Table 3). We tracked the power of tremor frequency oscillations as a function of varying the conductances. Relative to baseline at 1.0, tremor power changes were calculated at GABA conductance levels of 0, 0.5, 2, and 4. While GABA_A_ from TRN to VIM did not significantly affect tremor in either the extrinsic or intrinsic models (Fig. 5A–B), increases in TRN-TRN GABA_A_ significantly reduced tremor power in both models (Fig. 5C–D, Table 3).

**Figure 5:**
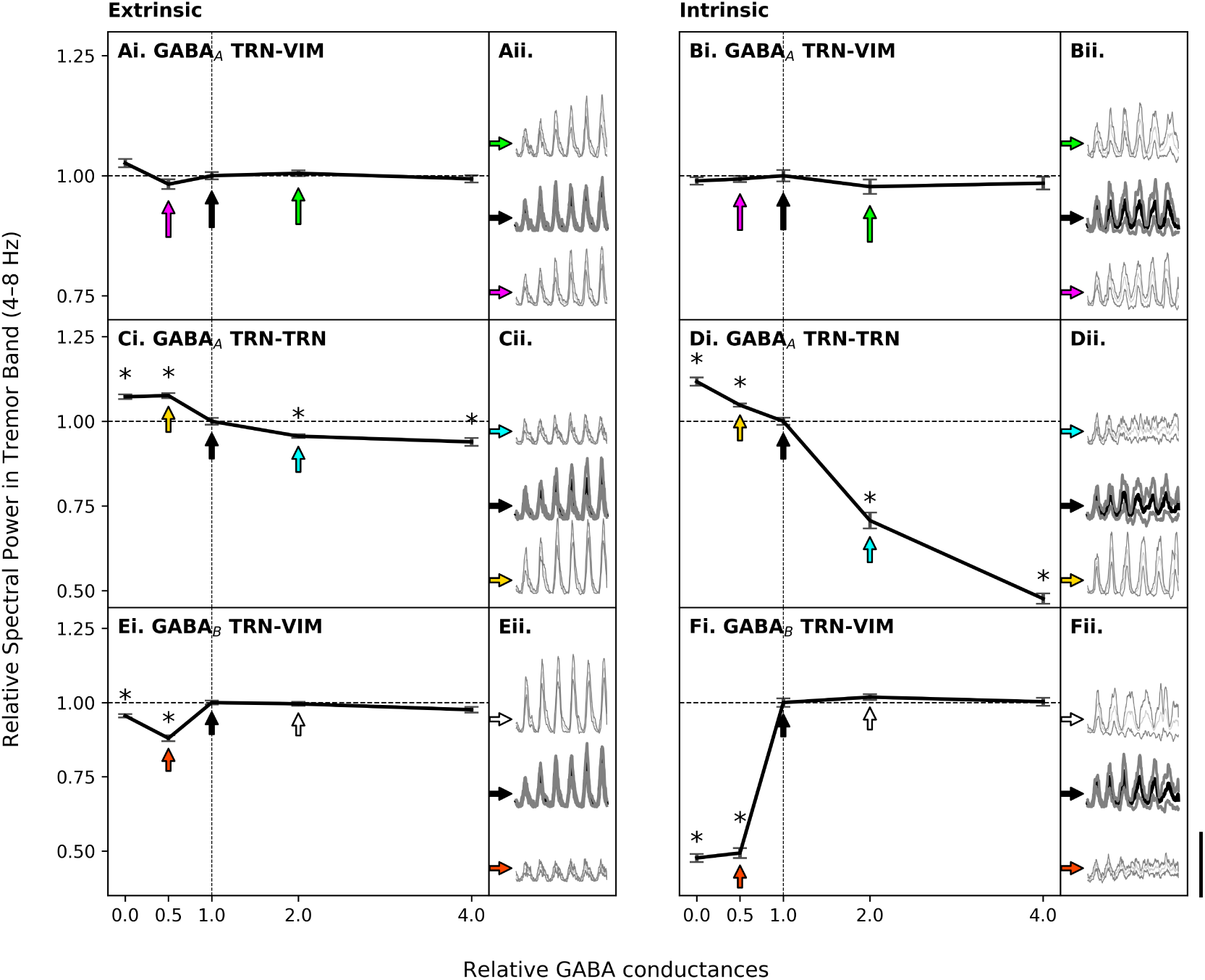
Tremor frequency power varied by GABA type in both models. Conductances of TRN-VIM GABA_A_, TRN-TRN GABA_A_, and TRN-VIM GABA_B_ were varied from baseline (normalized 1.0). Differences in tremor frequency power from baseline are plotted as a function of relative conductance. Colored arrows in i panels correspond to mean spike histograms in ii panels. Scale bar for all spike histograms is 25 sp/s. Values in Table 3. A–B. GABA_A_ from TRN-VIM. No changes affected tremor frequency power for either the extrinsic or intrinsic models. C–D. GABA_A_ from TRN-TRN. For both extrinsic and intrinsic networks, increases in conductance reduced tremor frequency power, and reductions in conductance increased tremor frequency power, suggesting that a global increase in GABA_A_-ergic tone may reduce tremor by affecting TRN-TRN. Total removal led to significantly greater increases in tremor frequency activity in the intrinsic compared to the extrinsic network. Increases at 2.0 and 4.0 led to significantly lower tremor in the intrinsic compared to the extrinsic network. E–F. GABA_B_ from TRN-VIM. For both extrinsic and intrinsic networks, reductions in conductance reduced tremor frequency power, and tremor frequency power values in the intrinsic network were significantly lower than in the extrinsic network.

**Table 3:**
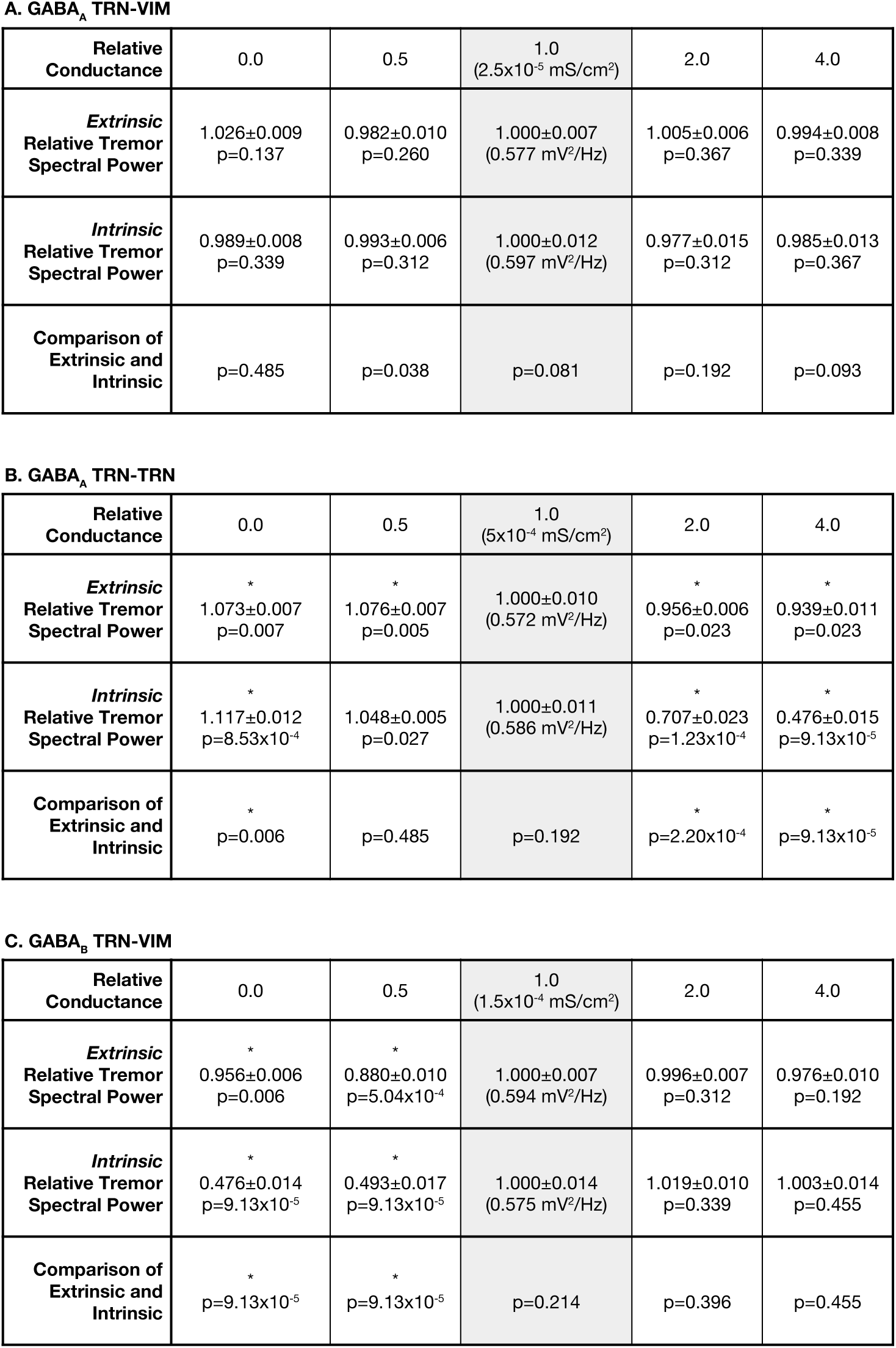
Modulation of tremor frequency power by GABA Conductance in models.

In both networks, reductions in TRN-TRN GABA_A_ increased tremor frequency activity relative to the respective baselines. In contrast, as the conductance was increased, in both models, significant reductions in tremor frequency activity were observed. Tremor power was directly compared at each conductance level (Table 3), and the TRN-TRN GABA_A_ conductance had a greater impact on the intrinsic versus extrinsic networks. With TRN-TRN GABA_A_ conductance removed, the intrinsic tremor frequency power was significantly greater than in the extrinsic network.

With increases in TRN-TRN GABA_A_, both the extrinsic and intrinsic networks had reduced tremor frequency activity. However, the tremor frequency activity in the intrinsic network was significantly lower than that of the extrinsic network at both 2.0 and 4.0 times the baseline conductance.

In the extrinsic model, the relatively smaller changes in tremor frequency activity due to changes in TRN-TRN GABA_A_ were due to the fact that an extrinsic drive was propagating tremor frequency activity in the network. The VIM-TRN network was more resilient to changes in these dynamics than the intrinsic network. The intrinsic network, having no external driver of tremor frequency activity, was significantly paced by the presence of internal GABA_A_ dynamics in the TRN. When this was disturbed by increases in TRN-TRN GABA_A_, the TRN reduced the ability for the VIM to maintain tremor dynamics.

In both the extrinsic and intrinsic models, increased GABA_A_ conductance within the TRN reduced tremor frequency activity in the VIM. These model results predict that precise upregulation of GABAergic activity specifically within the TRN will reduce tremor more effectively than global, thalamic GABAergic modulation.

### 3.4 Tremor frequency power decreased with reductions in GABA_B_ conductance in both models

Next, we analyzed modulation of GABA_B_ for fixed GABA_A_ values. Reductions in TRN-VIM GABA_B_ conductances, but not increases, significantly decreased tremor frequency power (Fig. 5E–F). In the extrinsic network (Fig. 5E), the power of TFOs dropped at both half the conductance and with complete removal. Decreases in TRN-VIM GABA_B_ conductance also reduced intrinsic power (Fig. 5F). Reductions in tremor power due to TRN-VIM GABA_B_ reductions were also significantly greater for the intrinsic model compared to the extrinsic model. Extrinsic tremor frequency activity, though paced by the tremor input, was still affected by the reduction of GABA_B_. However, intrinsic activity was strongly influenced by reductions of TRN-VIM GABA_B_ in the model, suggesting that treatments targeting GABA_B_ in the VIM-TRN circuit may help to reduce tremor, irrespective of the generating mechanism. At least one such drug, clozapine, may effectively reduce tremor in ET [8] through anatagonism of GABA_B_ [63].

Overall, both GABA_A_ increases within TRN and GABA_B_ reductions from TRN to VIM reduced tremor frequency activity in both the extrinsic and intrinsic networks. Additionally, reductions in GABA_A_ within the TRN led to increased tremor frequency activity in the VIM, suggesting that precision in GABA modulation is important in controlling the expression of tremor.

### 3.5 Intrinsic tremor frequency activity was further increased by tremor periodic input

It has been suggested that tremor frequency activity may be driven by cerebellum [15, 37, 52], and it is also possible that cortex plays a role [2, 57]. Yet it is not known exactly how tremor frequency activity interacts among different regions of this circuit and specifically how extrinsic influences affect the intrinsic oscillatory capacity of the thalamus. To account for the possibility of these interactions co-occurring, we simulated intrinsic oscillations combined with a simultaneous, extrinsic 6 Hz tremor frequency input. The simulated phasic drive consisted of 30 normally distributed AMPA-ergic inputs per cycle of the 6 Hz oscillation. As the unitary input strength was increased from 1×10^−4^−5×10^−3^ *μS*, tremor frequency power was significantly increased at 5×10^−4^ and 5×10^−3^ *μS* compared to the network with no input (Fig. 6 and Table 4A).

**Figure 6:**
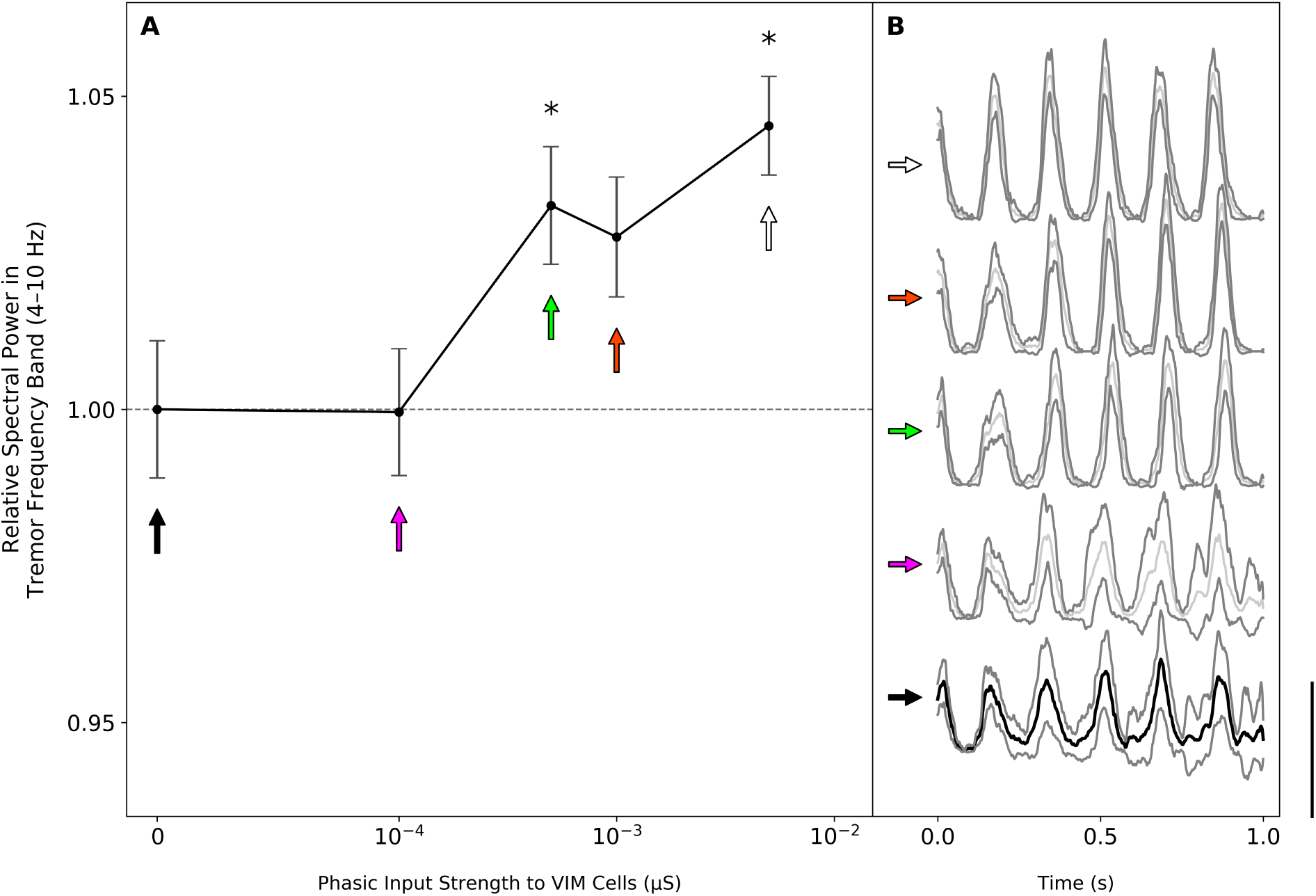
Intrinsic network tremor frequency power increased with tremor-phasic input. Phasic, 6 Hz synaptic inputs onto the VIM cells increased the tremor spectral power of intrinsically generated tremor oscillations. Values in Table 4A. A. Increased strengths of phasic input increased tremor activity with ongoing intrinsic oscillations. B. Mean spike histograms for simulations at each value of input. Arrows correspond to points in (A). Dark black line denotes mean intrinsic tremor in the absence of additional inputs. Gray lines are standard error. Scale bar is 25 sp/s.

**Table 4:** Modulation of tremor frequency power by synaptic inputs in models. A. Tremor-periodic inputs into the intrinsic network increased tremor. B. Non-tremor inputs increased tremor in intrinsic network but not extrinsic network.

**A. Tremor-periodic Input into Intrinsic Network**
| Unitary Input Strength (μS) | 0.00 | $1 \times 10^{-4}$ | $5 \times 10^{-4}$ | $1 \times 10^{-3}$ | $5 \times 10^{-2}$ |
| --- | --- | --- | --- | --- | --- |
| Tremor Frequency Power | $1.000 \pm 0.109$<br>($0.601 \pm 0.007$ mV<sup>2</sup>/Hz) | $1.000 \pm 0.010$<br>p=0.425 | $1.033 \pm 0.009$<br>p=0.023 | $1.028 \pm 0.010$<br>p=0.027 | $1.045 \pm 0.008$<br>p=0.001 |

**B. Poisson-distributed Input**
| Unitary Input Strength (μS) | 0.00 | $2.5 \times 10^{-5}$ | $5.0 \times 10^{-5}$ | $7.5 \times 10^{-5}$ | $1.0 \times 10^{-4}$ |
| --- | --- | --- | --- | --- | --- |
| Extrinsic Tremor Frequency Power | $1.000 \pm 0.012$<br>($0.586 \pm 0.007$ mV<sup>2</sup>/Hz) | $0.975 \pm 0.018$<br>0.136 | $0.948 \pm 0.014$<br>p=0.016 | $0.967 \pm 0.017$<br>p=0.052 | $0.906 \pm 0.015$<br>p=5.04x10<sup>-4</sup> |
| Intrinsic Tremor Frequency Power | $1.000 \pm 0.011$<br>($0.589 \pm 0.011$ mV<sup>2</sup>/Hz) | $1.035 \pm 0.008$<br>p=0.084 | $1.056 \pm 0.013$<br>p=0.015 | $1.052 \pm 0.014$<br>p=0.024 | $1.027 \pm 0.015$<br>p=0.132 |
| Comparison of Extrinsic and Intrinsic | p=0.299 | $2.17 \times 10^{-3}$ | $1.88 \times 10^{-4}$ | $1.10 \times 10^{-3}$ | $1.88 \times 10^{-4}$ |

These oscillations were strongly driven by the external input, which is seen in the mean fields in Fig. 6B. From baseline, as the strength of the external drive increased, the oscillations became more regular. The increases in tremor power suggested that the intrinsic activity of the thalamus can co-exist with external influences from areas such as deep cerebellar nuclei or the primary motor cortex.

### 3.6 Simulated non-tremor frequency activity reduced tremor power in the extrinsic network but increased tremor power in the intrinsic network

Cerebellar motor and non-motor activity is transmitted from deep cerebellar nuclei to the VIM, and primary motor cortex also projects to VIM [2, 57], so we simulated how different tremor mechanisms in VIM responded to non-tremor inputs that could arise from these sources. We simulated different strengths of a Poisson-distributed AMPA-ergic input with a rate parameter *λ* of 25 ms (see Methods) and found that the input significantly decreased tremor power in the extrinsic model and significantly increased tremor power in the intrinsic model (Fig. 7 and Table 4B).

**Figure 7:**
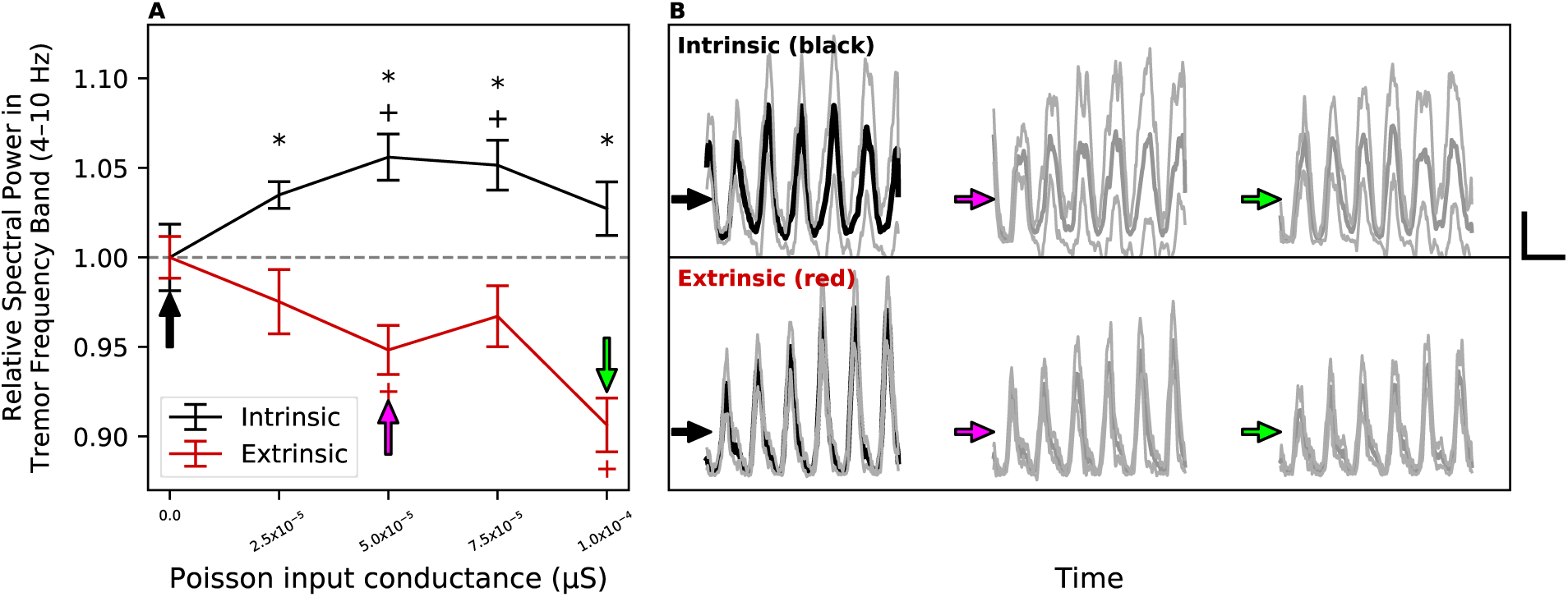
Non-tremor frequency inputs increased intrinsic but not extrinsic tremor frequency activity. An AMPA-ergic Poisson-distributed input was simulated as a model of non-tremor motor activity from cerebellum or cortex. Scale bars for B are 0.2 s and 5 sp/s. Values in Table 4B. A. The tremor frequency power relative to the baseline was tracked as the input conductance was increased for both the extrinsic (red) and intrinsic (black) model networks. As the input conductance increased, tremor frequency power in the extrinsic network was significantly reduced from baseline (red +), and the tremor frequency power in the intrinsic network was significantly increased from baseline (black +). For the same input, tremor frequency power was significantly greater in the intrinsic network compared to the extrinsic network (black asterisks). Values for the conductances and spectral power in Table 4. B. Mean VIM cell spike convolutions for the intrinsic and extrinsic networks are shown for 3 input strengths. The Poisson input led to increases in power for the intrinsic network. In the extrinsic network, the Poisson input interfered with the tremor-periodic input driving the extrinsic activity, reducing the overall amplitude. Colors corresponded to A.

In the extrinsic model, as the strength of the Poisson input was increased from 2.5×10^−5^−1×10^−4^ *μS*, the tremor power decreased from the baseline, significantly at 5×10^−5^ and 1×10^−4^ *μS*. Because the extrinsically driven tremor frequency activity was dependent upon the tremor-periodic input, an additional non-tremor input interfered, reducing the amplitude of tremor frequency activity in the network (Fig. 7Bi).

In contrast, in the intrinsic model, there was a significant increase in tremor concentration with an increase of Poisson conductance from 5.0×10^−5^−1×10^−4^ *μS*. As shown in Fig. 7Bii, though the amplitude of the tremor frequency activity did not increase, the shape of the activity was less sharp, due to the Poisson input broadly promoting periods of ongoing excitation in the intrinsic tremor frequency activity.

For each Poisson input conductance tested, the resulting tremor power in the extrinsic model was directly compared to that of the intrinsic model. In each case, the tremor power was significantly less in the extrinsic model than the intrinsic model. While Poisson inputs to the extrinsic network disrupted the tremor frequency activity, they facilitated tremor frequency activity in the intrinsic network, supporting the view that non-tremor cerebellar or cortical inputs into VIM would increase tremor frequency activity if dominated by intrinsic mechanisms, while the same inputs would reduce tremor frequency activity if it is driven by extrinsic mechanisms.

## 4 Discussion

We simulated a network of VIM cells and TRN cells that generated tremor frequency oscillations to understand how two distinct mechanisms might contribute to the generation of these oscillations and how they respond to non-tremor motor activity. The ultimate motivation of our study is to improve treatment strategies for ET, which rely on an understanding of the mechanisms of generation of tremor. “Extrinsic” tremor frequency activity was driven by tremor-periodic inputs from an external source like the dentate nucleus of the cerebellum. “Intrinsic” tremor frequency activity was generated by self-paced rhythmic activity that relied on GABA-ergic TRN inputs onto the cells in VIM and the T-type Ca^2+^ activity in both VIM and TRN cells. Both of these networks demonstrated tremor frequency field activity, but their mechanisms of generation and responses to non-tremor frequency activity were distinct, suggesting that potentially different treatment strategies may be necessary to alleviate different generators of tremor frequency activity. Though recent computational work has investigated various aspects of network level dynamics of tremor frequency activity in a mean-field sense [65], to our knowledge, this is the first time the VIM-TRN network with cellular biophysics has been simulated to investigate the cellular and network level origins of tremor periodic activity in ET.

### 4.1 Non-tremor cerebellar inputs may exacerbate tremor in VIM

The cerebellum is thought to contribute to motor planning and adaptation, and disruption of the cerebellothalamocortical pathway via lesion or deep brain stimulation (DBS) has been shown to impair these functions [9]. Here, we focused particularly on understanding how the membrane biophysics in VIM and TRN cells influenced larger-scale properties of the resulting network.

To understand effects of non-tremor frequency activity in ET [32], we simulated Poisson-distributed synaptic inputs from cerebellum or possibly cortex that are related to features such as motor adaptation. Our results showed that non-tremor inputs potentiated intrinsic tremor and depressed extrinsic tremor (Fig. 7), suggesting that activating inputs would increase tremor output if intrinsic oscillation are being generated within VIM, while extrinsically paced tremor in VIM would be reduced during high task demands. Indeed, ET patients typically present with an “intention” tremor, in which directed movement evokes tremor, consistent with the intrinsic model. Additionally, tremor frequency activity might increase during behaviors with prominent cerebellar contributions, such as motor learning, if intrinsic oscillations are being generated within VIM. Extrinsic activity would show reductions in tremor. It is possible that the movement-related activity from cerebellum gets transmitted as tremor frequency activity when any movement is made that is tremulous, consistent with both the intrinsic and extrinsic models. The hypothesized distinction between tremor frequency activity due to different inputs to VIM may be directly tested by recording from regions of VIM that exhibit tremor frequency activity in the absence of tremulous movement and recording changes in activity during a task parametrically varying motor intention.

DBS has also been suggested to reduce tremor by targeting cerebellothalamic fibers [30, 51], which is consistent with the view that tremor frequency activity is driven by cerebellar inputs. Yet the model results demonstrated that non-tremor frequency activity also increased tremor in the intrinsic model (Fig. 7), and this increase may be critical to the translation from tremor frequency neural activity to the expression of behavioral tremor. If tremor frequency activity in VIM must cross a threshold to drive behavioral tremor expression, then DBS affecting cerebellothalamic fibers may be eliminating non-tremor inputs to VIM, still resulting in reduced tremor output despite not explicitly blocking tremor frequency input activity. In a companion paper, we investigate the influence of symmetric, biphasic DBS patterns on both extrinsic and intrinsic generation of tremor frequency activity [33].

### 4.2 Broad effects of GABA modulation on tremor frequency activity in VIM

The contribution of GABA-ergic mechanisms to ET is well established in numerous studies [1, 4, 15, 44], though many are focused on the effects of GABA in the cerebellar system and less on the direct effects of GABA in the VIM, which our results suggest may be equally important to consider.

In some manifestations of ET, postmortem studies have demonstrated that Purkinje cells are atrophied and degenerated, resulting in a reduction in GABA-ergic tone [38], though this is disputed [49]. Cerebellar deficiencies in ET may play a role in at least some pathology seen in ET, which may be a more heterogeneous disease than commonly thought.

In the model studied here, to generate tremor frequency oscillations in the intrinsic model, the GABA_B_ strength was greater than that of GABA_A_. The model results suggested that TRN-VIM GABA_A_ did not have an effect on TFOs (Fig. 5), while both TRN-TRN GABA_A_ and TRN-VIM GABA_B_ affected tremor frequency activity in the VIM-TRN circuit. Increases in TRN-TRN GABA_A_ reduced TFOs in VIM, consistent with observations in tremor reduction with ethanol in alcohol-responsive forms of ET.

Within the VIM, the reductions in TRN-VIM GABA_A_ have been implicated in tremor frequency activity, but this was not observed in the present model (Fig. 5). Microinjections of the GABA_A_ agonist muscimol in VIM have been shown to attenuate tremor after several minutes [1]. It remains an important question how direct injections in or near TRN could affect tremor frequency activity. According to the model results, similar microinjections either directly in or closer to the TRN would result in more rapid cessation of tremor.

There is limited evidence that the effect of muscimol may not be selective to GABA_A_ but also affect presynaptic GABA_B_ [60]. If true in the VIM, this would provide support for the effects of GABA_B_ in the models. In the model, reductions in GABA_B_—similar to the possible effects of muscimol at presynaptic sites—resulted in reduced tremor generated either by extrinsic or intrinsic mechanisms (Fig. 5E–F).

In another study using positron emission tomography, an increased binding of ^11^C-Flumazenil at GABA_A_ receptors was observed in the cerebellum, premotor cortex, and the VIM in ET patients compared to controls, suggesting either increased upregulation of these receptors or a deficiency in normal binding [4]. While it may be possible that this finding was in part due to deficiencies in neighboring regions of TRN, further work on GABA_A_ and GABA_B_ deficiencies is necessary, to understand whether increases in TRN-TRN GABA_A_ and reductions in TRN-VIM GABA_B_ reduce tremor as predicted by the model.

### 4.3 T-type Ca^2+^ currents affected tremor frequency activity in VIM models

T-type Ca^2+^ channels have been long known to contribute to oscillatory behavior in the brain [10, 19], and they are present in areas that have been implicated in ET pathophysiology, such as the inferior olive and the dentate nucleus of the deep cerebellar nuclei. Also present in VIM, changes in the model T-type Ca^2+^ channels affected both extrinsic and intrinsic oscillations (Fig. 4). Increased T-type Ca^2+^ conductance reduced tremor frequency activity in both extrinsic and intrinsic models, though in different ways, while significant changes in frequency were observed for intrinsic but not extrinsic oscillations. In contrast, reductions in T-type Ca^2+^ currents resulted in significant increases in extrinsic tremor frequency activity and significant decreases in intrinsic tremor frequency activity.

Certain T-type Ca^2+^ antagonists, such as zonisamide [34, 40], have been attempted for treatment of ET [42], but the efficacy is not yet clear [43]. Zonisamide reduces T-type Ca^2+^, which our model suggests would lead to different effects in tremor power in VIM, depending on the prevalence of extrinsic versus intrinsic mechanisms (Fig. 4B and H). In the intrinsic model, reductions in T-type Ca^2+^ also led to a reduction in the tremor frequency (Fig. 4G).

In our network, increased T-type Ca^2+^ currents reduced tremor power in both networks and also increased the frequency of intrinsic tremor oscillations (Fig. 4). Despite the role of T-type Ca^2+^ currents in promoting oscillatory activity, in both models, too much of the current desynchronized the network activity and led to paradoxically less tremor frequency activity, providing a testable experimental prediction.

Here we used model neurons in which T-type Ca^2+^ was active only in dendritic compartments and affected output in other compartments in a non-linear manner. In a model of coupled single compartmental neurons with reciprocal inhibition that involved T-type Ca^2+^ channels, Shaikh *et al*. suggested that increased T-type Ca^2+^ currents reduced tremor frequencies and increased tremor power [53]. While additional work may be needed to rectify these results, the specific network topology may be critically important to understanding the outcomes of seemingly similar manipulations and suggests that pharmacological consequences may vary substantially depending on the anatomical site of action. Direct manipulation of T-type Ca^2+^ at relevant sites—such as the TRN, VIM, inferior olivary nucleus, and dentate nucleus—may help to delineate to what extent T-type Ca^2+^ disruptions at these sites can reduce tremor.

## Conclusions

We developed a model of VIM-TRN and simulated both extrinsic and intrinsic mechanisms of TFOs in the models to study their independent contribution to tremor generation. Improved understanding of these mechanistic differences is essential to devising effective, targeted treatment strategies. Our simulations suggested that these two distinct types of oscillations may co-exist and confer distinct properties that are important in understanding mechanisms of tremor. Manipulations in the model affected the magnitude and frequency of the tremor activity in VIM and could also affect the timing of expression with respect to behavior; understanding these properties may help delineate circuit mechanisms to give rise to better therapies. The model provides specific predictions that can be tested experimentally to understand the extent of intrinsically paced oscillations on tremor frequency activity in the thalamus and how these interact with external influences.

## Acknowledgements

This work was funded by NIMH 5T32MH019118-23, NIMH R01 MH106174 (SRJ), Brown University Dean’s Emerging Areas in New Science Award (SRJ and WFA), and the Doris Duke Clinician Scientist Development Award (WFA). The authors declare no competing financial interests.

